# Myeloid-Specific *Pck1* Deficiency Does Not Alter Aortic Root Atherosclerosis in Mice

**DOI:** 10.64898/2026.07.08.737280

**Authors:** Juying Han, Emmanuel Opoku, Jonathan D. Smith

## Abstract

**Background:** We previously performed a strain intercross between atherosclerosis resistant AKR *Apoe*^-/-^mice and atherosclerosis sensitive DBA/2 *Apoe*^-/-^ mice and identified the *Ath28* quantitative trait locus (QTL) on the distal end of chromosome 2. Congenic strain fine mapping identified the *Ath28*.*1* QTL atherosclerosis modifying subregion, encompassing 217 Kb, containing for only three protein-coding genes, *Zbp1, Pck1*, and *Pmepa1*, encoding respectively, Z-DNA binding protein 1, phosphoenolpyruvate carboxykinase 1, and prostate transmembrane protein androgen induced 1.

**Methods:** The effect of macrophage-specific knockout of *Pck1* (KO) was tested using the AAV2 transduced proprotein convertase subtilisin kexin type 9 (PCSK9) overexpression mouse model of hyperlipidemia and atherosclerosis.

**Results:** Unexpectedly, macrophage *Pck1* deficiency lowered body weight, liver weight, and HDL-cholesterol levels in both sexes, while total and non-HDL cholesterol levels were only decreased in male mice. Aortic root lesion area and necrotic lesion area were unchanged in KO mice of both sexes.

**Conclusion:** *Pck1* was not confirmed as an atherosclerosis modifier gene.

## Background

Atherosclerosis is a leading cause of heart disease that can lead to myocardial infarction and sudden cardiac death. Coronary artery disease (CAD) greatly increases in prevalence upon aging, and both genetic and environmental factors play large roles in its incidence. In both humans and in mouse models, it is clear that hyperlipidemia, specifically elevated non-HDL cholesterol, is required for atherogenesis. Monogenic forms of hyperlipidemia, such as familial hypercholesterolemia (FH) due to one defective copy of the LDL receptor gene, give rise to early onset coronary artery disease. In addition, genome wide association studies (GWAS) have identified hundreds of loci where common genetic variants are associated with coronary artery disease. Polygenic risk scores (PRS) calculated using thousands to millions of effect-size weighted common genetic variants have shown that individuals in the top 8% of the PRS have 3-fold greater prevalence of CAD compared to the remaining 92% of the population.^1^ Those in the top 8% of PRS are ∼ 20-times more common than FH in the general population and yet have CAD risk comparable to those with monogenic FH.

Considering the power of human GWAS in large population cohorts, one might ask why we bother to identify atherosclerosis modifier genes in mouse models? A GWAS study is just the beginning and due to linkage disequilibrium, further functional genomic studies are required to identify the causal gene, the causal variant, and the mechanism for the variant association with the human phenotype. Inbred mice are invaluable tools for research, as within each line all alleles are homozygous, and one can study environmental challenges without competing genetic variability. However, when comparing among different inbred lines, there is abundant genetic variability throughout their genomes. Researchers can take advantage of this variability in order to map loci that alter mouse phenotypes, which is performed via strain intercrossing or backcrossing followed by quantitative trait locus (QTL) analysis.^2^ Further fine mapping and candidate gene testing, including gene editing, can identify the causal gene and the mechanism of action, whether it be a protein coding change or a regulatory variant.^2,3^ This can identify modifier genes that do not contain common human protein changes or regulatory variants and thus may not reach the stringent threshold used in human GWAS studies. And, the identification of new modifier genes may lead to new pathogenesis pathways, new diagnostics, and novel therapeutic targets.

Using marker selected backcrossing, *Apoe* gene knockout (KO) was bred onto six inbred strains; and, among those the DBA/2 strain had the largest lesions and the AKR strain was one of several with the smallest lesions.^4^ An AKR *Apoe*^*-/-*^ x DBA/2 *Apoe*^-/-^ strain intercross identified several QTLs associated with aortic root lesion area, including the *Ath28* QTL on the distal end of chromosome 2.^5^ *Ath28* was confirmed in an independent AKR *Apoe*^*-/-*^ x DBA/2 *Apoe*^*-*/-^ strain intercross.^6^ Finally, *Apoe*^-/-^ congenic mice were created with the distal end of chromosome 2 derived from the DBA/2 background and the rest of the genome derived from the AKR background, which confirmed that the DBA/2 alleles on the end of chromosome 2 led to larger aortic root lesion areas.^7^ A series of additional congenic strains divided the *Ath28* QTL into three regions associated with lesion area, the smallest being the *Ath28*.*1* QTL region encompassing 217 Kb and harboring only three protein coding genes, *Zbp1, Pck1*, and *Pmepa1*.^7^ We previously tested *Zbp1* and found that its KO decreased atherosclerosis lesion and necrotic areas in C57BL/6 mice with hyperlipidemia induced by diet and weekly injections of an antisense oligonucleotide directed against the LDL receptor mRNA.^8^

Here we focus on the *Pck1* gene, encoding the cytosolic form of phosphoenolpyruvate carboxykinase, also known as PEPCK-C. It converts oxaloacetate and GTP into phosphoenolpyruvate plus GDP and CO_2_ with roles in hepatic and renal gluconeogenesis, as well as playing a role in cataplerosis (removal of excess oxaloacetate to promote the Kreb’s cycle) and glyceroneogenesis.^9^ In addition, human *PCK1* has been found to increase the generation of the methyl donor S-adenosylmethionine, promoting a specific histone 3 methylation (H3K9me3), which decreases hepatocellular carcinoma progression.^10^ The whole body KO of *Pck1* has a perinatal lethal phenotype, thus Cre-LoxP tissue specific knockout mice have been useful in studying the role of *Pck1* in various tissues.^11^

In the current study, we evaluated the effect of myeloid specific *Pck1* deficiency using hyperlipidemia induced by a one-time infusion of *PCSK9* encoding adeno associated virus (AAV) followed by feeding a western-type diet. KO mice of both sexes had significantly lower body weight and liver weight, as well as HDL cholesterol levels vs. their controls. We observed lower plasma total cholesterol and non-HDL cholesterol only in the male KO vs. controls. There were no significant effect of this KO on aortic root lesion areas in both female and male mice, vs. their controls.

## Methods

### Mouse model

All mouse procedures were approved by the Cleveland Clinic IACUC, and the use of recombinant adeno associated virus 9 (AAV9) was approved by the Cleveland Clinic Biosafety Committee along with the Institutional Animal Care and Use Committee. AAV9 injections into mice were performed in an ABSL2 mouse quarantine room. Macrophage specific *Pck1* KO mice on the C57BL/6 genetic background were created by crossing Pck1^flox/flox^ mice (reference ^11^) with LysM-Cre+/-transgenic mice (JAX catalog #004781), as previously described ^12^, with breeders kindly supplied by Colleen Croniger at Case Western Reserve University. Mice with Pck1^flox/flox^ LysM-Cre+/-genotype will be referred to as KO and mice with the Pck1^flox/flox^ LysM-Cre-/-genotype will be referred to as wildtype (WT) controls.

Recombinant AAV9 encoding gain of function D377Y mutation PSCK9 ^13^ was purchased from the University of Pennsylvania Vector Core, lot V8385R, and stored in small aliquots at -80°C. We performed a pilot study in C57BL/6 mice to determine the dose of the rAAV9-PCSK1 required to induce hyperlipidemia via i.p. injection followed by 4-weeks of feeding a high-fat (21.2% by weight) high-cholesterol (0.2% by weight) western-type diet (Envigo catalog # TD.88137). The final dose chosen was 1×10^12^ virus particles per mouse delivered in a final volume of 0.2 ml sterile saline. The atherosclerosis study protocol used male and female WT and KO mice at 8 weeks of age injected i.p. with the rAAV-PSCK9 followed by feeding the western-type diet for 16 weeks with sacrifice, after a 4-6 hour daytime fast, at 24 weeks of age. No anesthesia was used, and mice were euthanized by exposure to CO_2_, as approved by the Cleveland Clinic IACUC. After euthanasia, the chest was opened and blood was retrieved by cardiac puncture, followed by organ removal.

### Biochemical and histological assays

EDTA plasma was prepared by centrifugation of whole blood for lipoprotein and total triglyceride analyses. To determine HDL-cholesterol levels, 30 μl of plasma was mixed with 30 μl of KBr (density of 1.12 g/ml), placed into 0.2 ml tubes and centrifuged at 70,000 rpm for 16 hours in an S100-AT3 rotor (Thermo Fisher Scientific). The bottom 30 μl layer with a density > 1.063 was used to determine HDL-cholesterol. Plasma was diluted 10-fold with PBS for determination of total cholesterol (TC) levels. Both assays were run with technical duplicates, along with cholesterol standards, using the Cholesterol Liquicolor kit (StanBio Laboratory #1010-225) adapted for a 96 well plate format. Non-HDL cholesterol levels were calculated as TC – HDL-cholesterol. Mice with non-HDL cholesterol <400 mg/dl, most likely due to issues with the rAAV-PCSK9 injection, were removed from the study.

Quantitative assessment of atherosclerosis and necrotic core in the aortic root was performed as previously described.^14^ Lesion areas were quantified as the mean value in six sections at 80 μm intervals using Image Pro software (Media Cybernetics). The necrotic areas were defined as lesion areas lacking cell nuclei.

### Statistics

All statistics were performed using GraphPad Prism software (Version 10.1.2). Female and male mice were analyzed separately, as recommended.^15^. All data were analyzed using the non-parametric two-tailed Mann-Whitney test unless stated otherwise.

## Results

### Induction of hyperlipidemia in control and myeloid-specific Pck1 KO mice

Hyperlipidemia in *Pck1*^flox/flox^-LysM-Cre+ (KO) and their littermate *Pck1*^flox/flox^ WT controls was induced, without the need for crossing to *Apoe*- or *Ldlr*-deficiency, by using a previously described PCSK9 encoding AAV and feeding a western-type diet.^16^ Mice were injected with the AAV at 8 weeks of age and then fed the high-fat high-cholesterol diet for another 16 weeks, as described in detail in the Methods section. Although these KO vs. control mice were previously reported to have no difference in immune cell frequencies, that study did not comment on body or organ weights.^12^ In the current study, upon sacrifice at 24 weeks of age, body weights were significantly reduced in the KO vs. WT control mice in both females (31.1% median decrease, p=0.0002) and males (26.2% median decrease, p<0.0001) (Figure 1A). Liver weights were also significantly reduced in the KO vs control mice in both females (30.2% median decrease, p=0.0008) and males (40.4% median decrease, p<0.0001) (Figure 1B). Spleen weights were not significantly different in the KO mice vs. controls for both sexes (Figure 1C).

**Figure 1.**
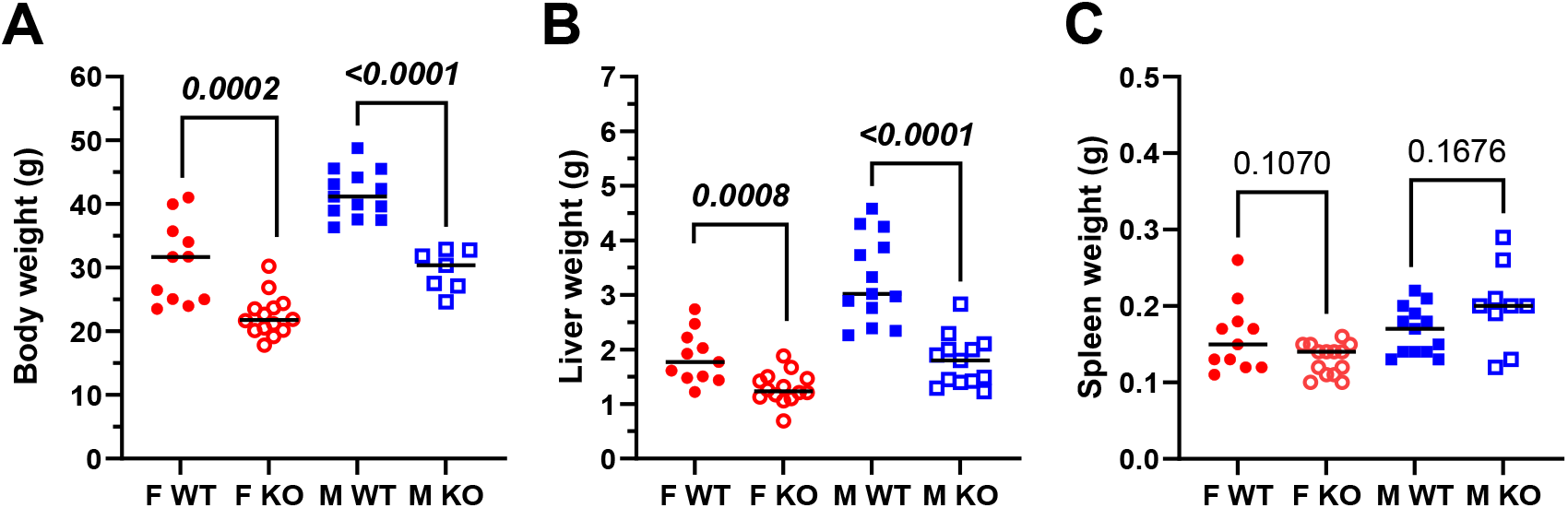
Macrophage *Pck1* KO alters body and organ weights. (**A**). Body weights. (**B**) Liver wet weights. (**C**) Spleen wet weights. Female wildtype controls, F WT, filled red circles; female KO, F KO, open red circles; male wildtype controls, M WT, blue filled squares; male KO, M KO, open blue squares. P-values within each sex determined by non-parametric two tailed Mann Whitney test, black lines denote the median values.

Plasma total cholesterol levels were unchanged in the female, but were significantly reduced in the male KO vs. WT mice (32.7 %median decrease, p=0.0033, Figure 2A). HDL-C values were significantly reduced in both the female (59.9%median decrease, p=0.0001) and male KO vs WT mice (52.4%median decrease, p<0.0001, Figure 2B). Non-HDL cholesterol, similar to total cholesterol, was only significantly reduced in the male mice (39.8% median decrease, p=0.0071, Figure 2C). Plasma triglyceride levels were unchanged in KO vs. WT mice of both sexes (Figure 2D).

**Figure 2.**
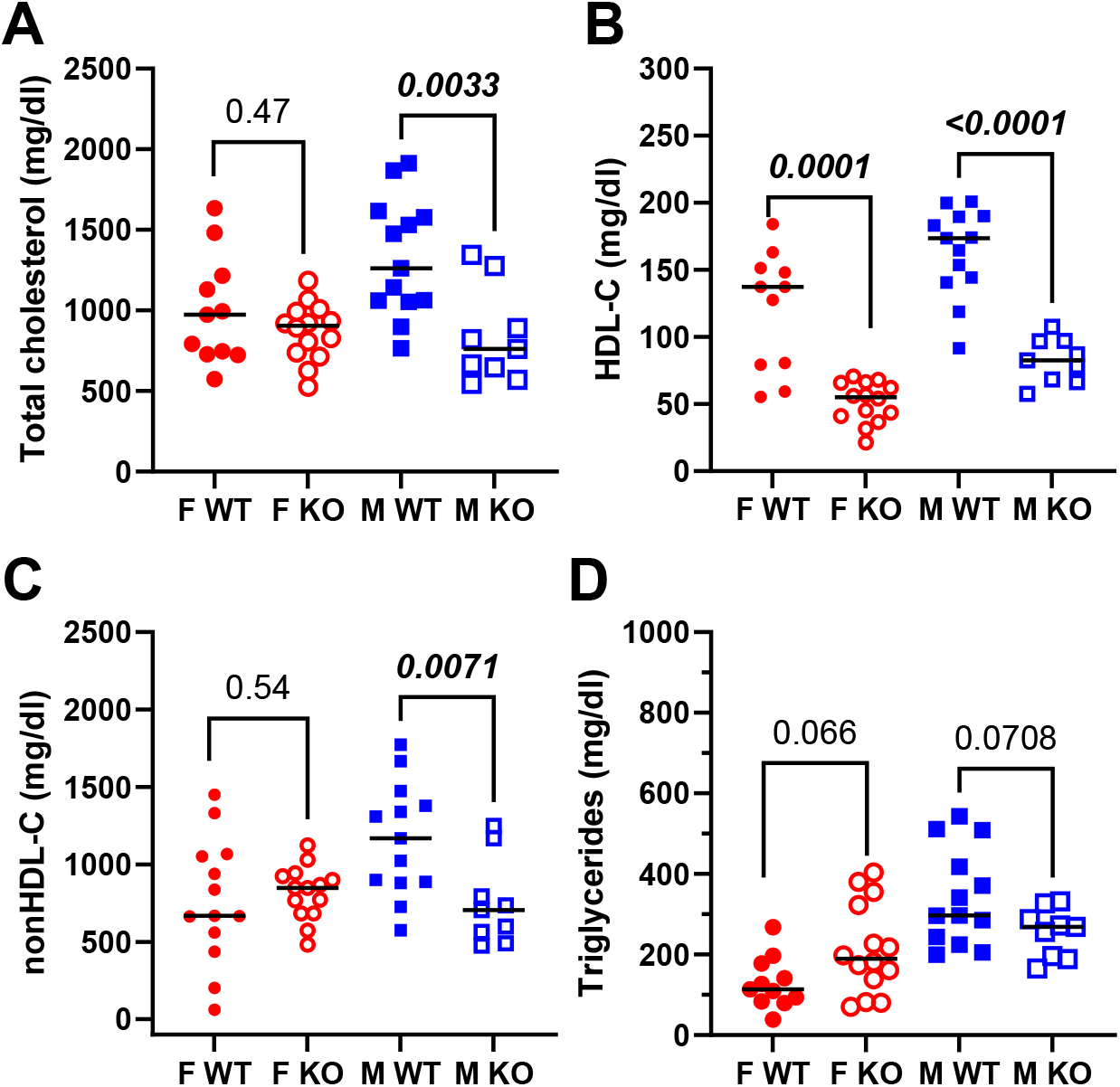
Macrophage *Pck1* KO effect on plasma lipid levels. (**A**). Total cholesterol. (**B**) HDL cholesterol (HDL-C). (**C**) Non-HDL cholesterol (nonHDL-C). (**D**) Triglycerides. Symbols the same as in Figure 1. P-values within each sex determined by non-parametric two tailed Mann Whitney test, black lines denote the median values.

### Aortic root atherosclerosis

Aortic root lesion areas and necrotic areas were assessed in mice at 24 weeks of age. Lesion areas were quite variable even within each genotype and sex, with the coefficient of variation (S.D./mean) ranging from 40 to 68% (Figure 3A). Lesion area in KO vs. WT mice were not significantly different in either sex, although the median lesion areas trended 14.7% and 26.1% larger in the KO vs. WT mice. In addition, there were no significant effects of the KO on necrotic lesion area in both sexes, with the coefficient of variations ranging from 81to 115% (Figure 3B). To assess whether the variability in lesion area was associated with non-HDL cholesterol levels, linear regression analyses were performed. There were no significant correlations between lesion area and non-HDL cholesterol within each of the four groups based on their sex and genotype.

**Figure 3.**
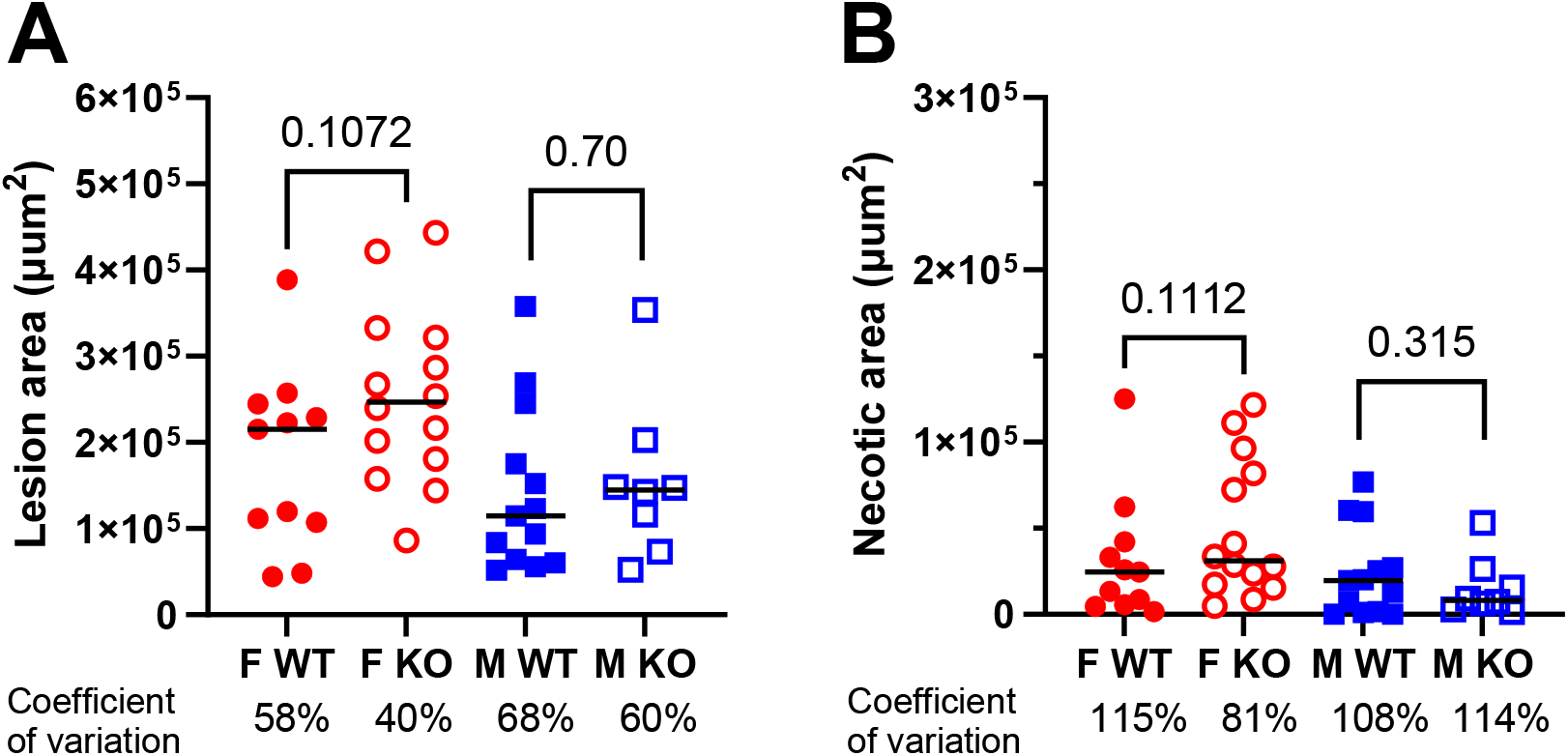
Aortic root atherosclerosis. (**A**). Lesion areas. (**B**) Necrotic lesion areas. Symbols the same as in Figure 1. The coefficient of variation is displayed under each sex and genotype group. P-values within each sex determined by non-parametric two tailed Mann Whitney test, black lines denote the median values.

## Discussion

In the current study we assessed *Pck1*, one of three protein coding genes within the *Ath28*.*1* QTL interval, as an atherosclerosis modifier gene. Although *Pck1* is expressed highly in hepatocytes, single cell RNAseq of mouse liver shows its expression in all cell types including macrophages.^17,18^ Similarly, single cell RNAseq of mouse aorta also shows *Pck1* expression in all cell types, including macrophages and monocytes, although expression was highest in endothelial cells.^18,19^ Since whole body *Pck1* KO is leads to perinatal lethality,^11^ we assessed its KO in myeloid cells based on use of *Pck1*^flox/flox^ and LysM Cre transgenic mice. Myeloid-specific *Pck1* KO mice are viable and their macrophages, studied *in vitro*, have increased glycolysis and decreased *de novo* lipogenesis, as well as increased proinflammatory cytokine production in response to lipopolysaccharide treatment.^12^ Many studies have provided an abundance of evidence for the roles of macrophages, lymphocytes, vascular smooth muscle cells, and endothelial cells in modifying atherosclerosis severity in various hyperlipidemic mouse models.^20^ However, macrophages seem to be particularly crucial for atherogenesis in mice, as several independent studies of hyperlipidemic mice with mutations in *Csf1* gene, also known as MCSF for macrophage colony stimulating factor, have 10 to 100-fold smaller lesions than their respective control mice, despite having even higher plasma levels of non-HDL cholesterol.^21–23^ Thus, we decided to test for the effect of *Pck1* deficiency in myeloid cells on atherosclerosis. The predominant myeloid cells in the aorta are macrophages and monocytes; and, although there is limited neutrophil accumulation in atherosclerosis due to their short lifespan, neutrophils may still play an important role in atherogenesis and recruitment of monocytes.^24^

We were surprised to observe that the myeloid KO of *Pck1* decreased body weight and liver weight of the mice fed the western-type diet. We don’t know the reason for this, and it is beyond the scope of our study to investigate this; however, myeloid specific knockout of *Asxl2* also decreased bodyweight in mice fed a high fat diet.^25^ High fat diets can induce adipose tissue macrophages to accumulate and acquire a pro-inflammatory phenotype, and the *Asxl2* KO macrophages had reduced IL-1β and TNFα secretion after inflammasome activation.^25^ However, *Pck1* KO macrophages have increased mRNA expression of IL-1β and TNFα in response to LPS, opposite of what was observed in the *Asxl2* KO macrophages.^25^ Thus, this enigma will require additional studies to unravel.

We were also surprised to see decreased plasma total and non-HDL cholesterol in the male KO mice, and decreased HDL cholesterol in both female and male KO mice. It has previously been shown that the response to recombinant AAV-PCSK9 i.v. treatment is more pronounced in male vs. female mice fed a chow diet, with higher total cholesterol in the male mice, which is due in part to a higher hepatic level of the recombinant AAV in males vs. females.^26^ In fact, our study also found a trend towards higher total cholesterol levels and significantly more non-HDL cholesterol levels (p=0.034 by Kruskal Wallis non-parametric ANOVA with Dunn’s multiple comparison posttest) in male vs. female mice that were WT for *Pck1* (Figure 2A, C). However, in the myeloid specific *Pck1* KO, the levels of total and non-HDL cholesterol were similar in the male and female mice, suggesting that *Pck1* KO may have led to equal hepatic levels of recombinant AAV levels in the male and female mice. We attribute similar decreases in HDL cholesterol in the female and male *Pck1* KO mice due to the decreased liver weight in both sexes, as tissue specific *Abca1* KO mice have shown that liver ABCA1 contributes more to HDL biogenesis than other organs.^27^

Returning to the *Ath28*.*1* QTL interval with its three protein coding genes^7^, we now demonstrate that KO of myeloid expression of *Pck1* had no significant effect on aortic root atherosclerosis in both female and male mice. As male mice had lower levels of plasma non-HDL cholesterol, this may have obscured any potential effect of *Pck1* KO to increase atherosclerosis. However, we are confident of our finding in female mice, as *Pck1* KO did not alter plasma non-HDL cholesterol in the female mice. However, one limitation of our study is that we only evaluated the effect of *Pck1* expression in myeloid cells and we cannot rule out that *Pck1* expression in endothelial or smooth muscle cells could modify atherosclerosis. There is still one untested atherosclerosis candidate gene remaining in the *Ath28*.*1* interval, *Pmepa1*, encoding the prostate transmembrane androgen induced 1 protein, which may be misleading, as the human homolog of this gene is expressed highest in the aorta and tibial artery.^28^

## Conclusion

Lack of Pck1 expression in myeloid cells has no significant effect on aortic root atherosclerosis in mice made hyperlipidemic by overexpression of PCSK9 and feeding a western type diet. Unexpectedly, this KO led to lower bodyweight, liver weight, and HDL cholesterol levels.

## Abbreviations

AAV: adeno associated virus
*Apoe*: apolipoprotein E gene
CAD: coronary artery disease
FH: 
familial hypercholesterolemia
GWAS: genome wide association study
KO: knockout
*Pck1*: phosphoenolpyruvate carboxykinase 1 gene
PCSK9: Proprotein convertase subtilisin kexin type 9
*Pmepa1*: prostate transmembrane protein androgen induced 1 gene
PRS: polygenic risk score
QTL: quantitative trait locus
WT: wild type
*Zbp1*: Z-DNA binding protein 1 gene

## Declarations

### Ethics approval and consent to participate

All studies were approved by the Cleveland Clinic Biosafety and Institutional Animal Care and Use Committee. The mice used were derived from breeders supplied by Dr. Colleen Croniger at Case Western Reserve University, and Dr. Croniger gave consent for us to use these mice for the current studies.

### Consent for publication

All authors provided consent for publication.

### Authors’ contributions

J.H. performed studies, analyzed the results, wrote parts of the manuscript, and edited the manuscript. E.O. performed studies, quantified outcomes, and wrote part of the manuscript. J.D.S. conceived and planned the study, obtained funding, supervised the research team, analyzed the results, wrote parts of the manuscript, and edited the manuscript.

